# Bespoke sustainable 3D-printed labware for enhanced handling and standardization of tumor spheroid migration and invasion assays

**DOI:** 10.64898/2026.07.28.741193

**Authors:** Tobias Butelmann, Tobias Nicolaisen, V. Prasad Shastri

## Abstract

Three-dimensional (3D) cell culture models, particularly multicellular tumor spheroids, have become essential tools for studying cancer biology, drug screening, and preclinical testing due to their ability to mimic physiological tumor microenvironments. However, traditional invasion assays, such as Boyden-chamber- or Transwell-based systems, often suffer from variability introduced by spheroid handling and transfer, compromising data reproducibility. Here, we present a novel, 3D-printed migration and invasion platform —the MQ_m_-sert— designed to standardize and streamline spheroid-based invasion assays while maintaining spheroid integrity. Fabricated via fused filament fabrication using biobased polylactic acid, the MQ_m_-sert integrates a hanging-drop spheroid culture system (MQ_m_-sert) with a membrane-based invasion chamber (M-sert), eliminating the need for disruptive spheroid transfer steps. Using synthetic tumor environment mimics (STEMs) composed of breast cancer cells (MDA-MB-231 and MCF7), mesenchymal stromal cells (MSCs), and human pulmonary microvascular endothelial cells (HPMECs), we quantified invasion dynamics and cellular interactions. This innovation significantly reduces experimental variability, as demonstrated by lower variance in invasive cell mass dimensions and cell density compared to conventional workflows. Beyond biological insights, the platform aligns with sustainability goals by leveraging cost-effective, open-source 3D printing, reducing reliance on commercial labware, and addressing key challenges in 3D cell culture standardization.

## Introduction

Three-dimensional (3D) cell culture (CC) has already surpassed traditional adherent, two-dimensional (2D) CC and is widely adopted nowadays.^1^ This is achieved for 3DCC by providing physiological traits and properties, including gradients of different kinds, multiple cell types, spatial organization and differentiation, as well as the possible involvement of materials for mechanical stimuli and biological cues, which cannot be sufficiently replicated by 2DCC.^2^ Different methods for 3DCC exist, which can be divided into scaffold-free and scaffold-based techniques, producing 3D structures solely with cells or, with the help of an extracellular matrix (ECM) mimicking material.^3^ Among scaffold-free techniques, the hanging-drop method is well-established for cultivating cellular spheroids, is the mainstay of cancer biology research, and has propelled many important discoveries.^4–6^

In addition to serving as in vitro models that mimic tumor traits, such as cellular organization and a hypoxic/necrotic core, these models serve as a bridge to animal models, especially for drug screening and pre-clinical testing, and serve to reduce animal numbers for pre-clinical studies.^7, 8^ The refinement and robustness of tumor spheroid models have become even more relevant in light of recent EU and FDA mandates to phase out and replace animal testing with 3DCC for chemical, drug, and antibody testing.^9, 10^

A typical workflow for spheroid preparation using the hanging drop method is shown in Figure 1, and it involves two essential steps: harvesting and transfer for downstream processing. Spheroid preparation involves pipetting and additional centrifugation to harvest them. These manipulations impose mechanical strain on the cells and introduce user-dependent errors, such as loss of spheroids during transfer, loss of spheroid integrity during handling, and potential cell activation.^11^

**Figure 1:**
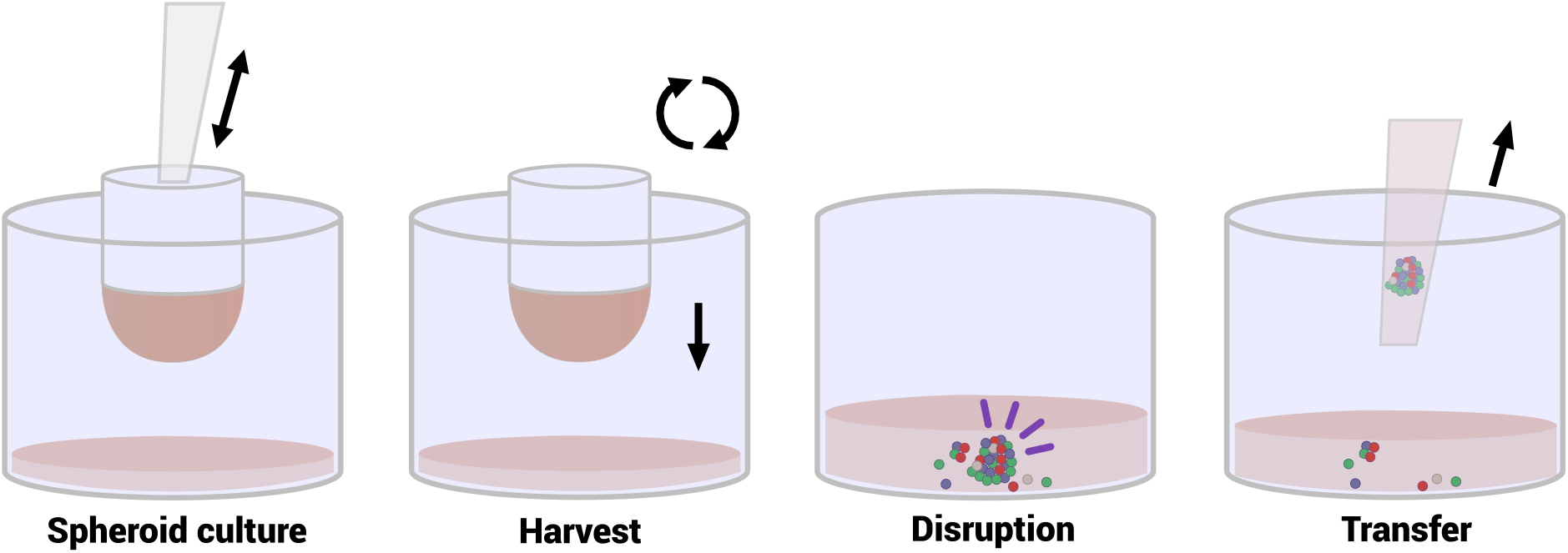
Traditional spheroid preparation by the hanging-drop method. It involves spheroid culture followed by two steps: harvest and transfer, which contribute to the disruption of spheroids.

For standardization, it is therefore important to limit and/or minimize human intervention and move towards automation, while adopting a cost-effective and sustainable workflow, to address the aforementioned EC and FDA roadmap.^12^ Standardization becomes even more consequential when spheroid culture systems are used to gather quantitative information on cellular processes related to metastatic potential, such as migration or invasion. Such assays require additional handling procedures, i.e., isolation of the spheroid (Figure 1), and transfer onto a membrane in a Boyden chamber or transwell assay. This step is a delicate procedure that introduces operator-dependent variables that can be difficult to mitigate and account for.^13, 14^ Based on these considerations, this study aimed to develop a system that improved workflow with less operator input and more standardizable practices, while incorporating sustainability elements.

In our previous work, we reported the development of inserts - Q-serts - for hanging-drop spheroid culture. The Q-serts featured a modular design with four hanging-drop chambers arranged in a four-leaf clover configuration. These chambers could be snapped into a standard 96-well cell culture plate and were fabricated from poly(L-lactic acid) (PLA), thus reducing waste by optimizing material use and enabling recycling of spent inserts. Additionally, the hanging-drop chamber design enabled automated spheroid handling using conventional liquid-handling robots, minimizing manual intervention.^15^ To ensure wide adoption and promote sustainable, economical fabrication of labware “on-demand”^16–20^, the Q-serts were fabricated using fused filament fabrication (FFF). Using the synthetic tumor environment mimic (STEM) spheroid system developed by our laboratory, which emulates cellular heterogeneity and stromal/vascular tumor compartments found in solid tumors or epithelial origin^21, 22^, the suitability of Q-serts for spheroid preparation was also validated.^15^

Building on this aforementioned work, this study introduces an innovative design that integrates spheroid preparation using the hanging-drop method with migration/invasion assays, enabling easy transfer of spheroids from hanging-drop culture to the chamber membrane. This system, termed MQ_m_-sert, consists of two parts. (1) An insert with a membrane (M-sert) for regular migration/invasion assays, and (2) a Q_m_-sert for hanging-drop culture of spheroids that is compatible with a single 96-well format. In contrast to the traditional two-step workflow involving spheroid isolation and transfer to the membrane (**Figure 1**), the MQ_m_-sert streamlines these two steps, eliminating errors due to the transfer step and yielding experimental outcomes that are highly reproducible and lack the deviations observed in the conventional workflow scenario.

## Materials and Methods

### Design, fused filament fabrication, and post-processing

Q-serts, M-serts, and Q_m_-serts were designed using Inventor Professional 2022 (Autodesk, USA) and exported as .stl files with a built-in feature. For exporting rendered images, the ray tracing option in the Inventor software was used with high lighting and material accuracy (processed until fine). Falcon cell culture inserts (Ref. No. 353097, Corning Inc., USA) were chosen as a reference to ensure comparison of M-serts with commercial products. A binary G-code was generated in PrusaSlicer 2.71 (Prusa, Czechia) with the following settings: layer height of 0.2 mm, printing speed of 12.5 mm/s (1^st^ layer 37.5 mm/s), a fan speed of 100 % (1^st^ layer 50 %), 15 % infill, a heated print bed at 60 °C and a temperature setting of 195 °C (1^st^ layer 210 °C). Polylactic acid filament (1.85 mm; Polymaker) was used to print parts on an MK4 (Prusa, Czechia) equipped with a 0.4 mm brass nozzle. The printed parts were checked for remaining filament strings, which were removed if necessary. Afterwards, they were rinsed in dH_2_O in an ultrasound bath (3 x 5 min) and blown dry with a compressed air gun.

### Device fabrication and sterilization

Following computer-aided design and FFF-based production, the membrane devices (M-serts) were processed and assembled as follows. An ipCELLCULTURE track-etched membrane (PET, 8 µm pore size, 6 × 10^4^ pores cm^-2^, it4ip, Belgium) was bonded to the printed scaffold with an epoxy resin mixture (PRO 2C, EPODEX, Germany) for 48 h at RT, and subsequently cured at 42 °C for 6 h. To test conformity and absence of leakiness around the seal in the fabricated devices, a 200 µm PET film devoid of pores (Volvic, France) was used instead of the track-etched membrane. Before cell culture, the parts were submerged in 70 % ethanol, sterilized under UV light (Puritec HNS 6W G5, Osram) for 30 min, and finally dried in the laminar flow cabinet after removing the ethanol. The devices were used within 30 days after fabrication.

### Cell culture

Cells were cultured at 37 °C under 5 % CO_2_ at ∼90 % humidity in their respective medium listed in Table 1. Cells were washed with DPBS and detached by trypsinization (0.05 % trypsin / 0.02 % EDTA) when they reached 70-80 % confluency. Treated cells assumed a rounded morphology and were then washed with DMEM supplemented with FBS to neutralize trypsin. Subsequently, cells were centrifuged, the culture medium removed, and the cells were suspended in fresh medium for another passage or taken to form spheroids.

**Table 1:** Cultured cells, cell lines and their culture media. P/S: penicillin/streptomycin. FGF2: fibroblast growth factor-2.

| Cell type, or line | Culture medium |
| --- | --- |
| Mouse fibroblasts NIH/3T3 (NIH3T3) | DMEM, 10 % FBS, 1 % P/S |
| Human lung adenocarcinoma cells (A549) | DMEM, 10 % FBS, 1 % P/S |
| Mouse hepatoma-derived cells (Hepa1-6) | DMEM, 10 % FBS, 1 % P/S |
| Human marrow-derived mesenchymal stroma cells E2Crimson-labelled (MSC E2C) | α-MEM, 10 % FBS, 1 % P/S, 5 ng/mL FGF2 |
| MDA-MB-231 blue fluorescent protein-labelled cells (MDA-MB-231 BFP) | DMEM, 10 % FBS, 1 % P/S |
| MCF7 green fluorescent protein-labelled cells (MCF7 GFP) | DMEM, 10 % FBS, 1 % P/S |
| Human pulmonary microvascular endothelial cells (HPMECs) | Endopan MV (P04-00200, Pan-Biotech GmbH, Germany) |

NIH3T3, MDA-MB-231, as well as MCF7 cell lines were procured from the toolbox of BLOSS/CIBSS (Center for Biological Signaling Studies, University of Freiburg, Germany). HPMECs were purchased from PromoCell (Cat. No. C-12281, Lot 480Z005, Heidelberg,

NIH3T3, MDA-MB-231, as well as MCF7 cell lines were procured from the toolbox of BIOSS/CIBSS (Center for Biological Signaling Studies, University of Freiburg, Germany). HPMECs were purchased from PromoCell (Cat. No. C-12281, Lot 480Z005, Heidelberg, Germany). MSCs were kindly provided by Prof. Andrea Barbero and were obtained from patients under consent per the regulations of the local ethical committee (University Hospital Basel; Ref No: 78/07).

### Lentiviral transduction

Lentiviral particles containing BFP: pLVX-mTagBFP2-P2A-Puro, E2 Crimson: pLVX-E2-Crimson-P2A-Puro (BIOSS Toolbox, University of Freiburg, Germany), or GFP: pGIPZ-turboGFP-IRES-PuroR (RHS4346, Horizon Discovery, United Kingdom) were produced in HEK293 cells by mixing transgene vector and packaging vectors (pCMVdR8.74 (packaging plasmid; Addgene, Plasmid #22036) and pMD2.G (envelope plasmid, Addgene, Plasmid #12259)) using branched polyethyleneimine (bPEI) (M_W_ = 25 kDa, Sigma Aldrich, Germany) as the transfection reagent. For transfection, 5 μg of DNA at a ratio of 4:3:1 (transgene: packaging: envelope plasmids) was diluted in 250 μL of Opti-MEM (Invitrogen, Thermo Fisher, USA), and 11.25 µL of bPEI (1 mg mL^-1^) was added and incubated for 25 min at room temperature before transferring to HEK293 cells. 16 h after transfection, the medium was replaced with a cell-specific culture medium, and the media containing lentiviral particles were collected after 24 and 48 h and filtered through a sterile 0.20 µm syringe filter (Millipore, Germany) to infect target cells. Three days after transduction, infected cells were selected using 2 mg mL^-1^ puromycin (Sigma-Aldrich, USA) in the respective culture medium.

### Spheroid culture

Sterilized Q-serts or Q_m_-serts were clipped into a 96-well plate for hanging drop cell culture. For each drop, 35 µL of a respective cell suspension was pipetted. Single-cell type spheroids consisted of 20,000 Hepa1-6 or A549 cells. A standard STEM spheroid consisted of a mixture of HPMECs, MSC E2C, and MDA-MB-231 BFP in 3:2:5 ratio for a total cell number of ∼22k cells per drop. The ratios were adjusted accordingly as indicated by adding MCF7 GFP. Plates were sealed with Parafilm (Amcor, Switzerland) to avoid excess evaporation. Spheroids were allowed to settle for 2 days, and an initial medium change of 5 µL (removal) and 7.5 µL (addition) was carried out, after which a daily medium change of 5 µL (removal) and 6 µL (addition) was applied. Spheroids from Q-serts were harvested using a cell culture centrifuge with a short pulse until 200 x g. Q_m_-serts were directly transferred into the M-sert, establishing an MQ_m_-sert.

### Mycoplasma test

In a 24-well plate, 500 µL DMEM supplemented with 10% FBS and 1% P/S was added per well, and a sterilized M-sert was attached to the well, after which the M-sert chamber was filled with 200 µL of medium. Samples were incubated for 72 h. After incubation, 500 μL of cell culture supernatant from a sample was transferred to a microcentrifuge tube. The sample was boiled at 95 °C for 10 min, then centrifuged for 5 s at 15,700 x g, and 150 µL of the supernatant was transferred to a fresh test tube for analysis by Eurofins Genomics (Ebersberg, Germany).

### MTT assay

In a 24-well plate, 500 µL DMEM 10 % FBS 1% P/S was added to a well, a sterilized M-sert was attached to the well, and the M-sert chamber was filled with 200 µL of medium, so that any cytotoxic substance could diffuse into the medium (Medium 1). As a control, 1.2 mL of medium was incubated in a 6-well. Five wells were prepared, and after 72 h, the medium was collected. The MTT assay was performed in a 96-well plate format. NIH3T3 cells were seeded at 10,000 cells per well in 100 µL medium. Cells were incubated overnight, then the medium was aspirated, and 100 µL Medium 1 was added for 24 – 48 h. Each of the five samples was split into three technical replicates, and 50 µL MTT solution (1 mg/mL) was added to each sample and incubated for 2 h. After carefully removing the MTT solution, 100 µL isopropanol was added, and the well plate was placed on a rotary shaker at 300 rpm for 15 min. Absorbance was measured using a BioTek Synergy HTX well-plate reader (BioTek Instruments, USA) at 570 nm.

### Transmembrane migration and invasion assays

#### Migration assay for single cell suspension

For migration studies of single cell suspensions, M-serts attached to 24-well culture plates (Ref. No. 83.3922, Sarstedt, Germany) were compared with Falcon cell culture inserts (Ref. No. 353097, Corning Inc., USA) attached to a 24-well companion plate (Ref. No. 353504, Corning Inc., USA) with an 8 µm pore size membrane at 6 x 10^4^ pores cm^-2^. Wells were filled with 500 µL DMEM (10 % FBS, 1 % P/S) per well, and the membrane of the respective cell culture insert was seeded with cells suspended in 150 µL serum-free medium at a density of 15,000 cells per membrane. The insert was placed in a pre-filled well and incubated for 24 h, and precautions were taken to avoid the entrapment of air bubbles under the membrane.

#### Invasion assay using single-cell suspensions spheroids

For an invasion assay, M-sert membranes were coated with 100 µL of Matrigel (300 µg/mL protein content, growth factor reduced, Cat. No. 354230, Corning Inc. USA) for 2 hours at 37 °C, 5 % CO_2,_ and ∼90 % humidity. The supernatant was carefully aspirated afterwards. For single-cell suspensions, 15,000 cells in 150 µL serum-free medium were seeded onto a membrane. Spheroids were prepared as described and cultured until day 7 to be transferred to the M-sert using two distinct workflows.

#### Traditional spheroid harvest (Q-sert)

Spheroids from Q-serts were harvested using a cell culture centrifuge with a short pulse at 200 x g, after which the Q-serts were removed using a pair of tweezers. Individual spheroids were carefully aspirated with minimal medium carry-over and dispensed in 200 µL fresh serum-free DMEM. After spheroid collection, 150 µL, including the spheroid, was carefully pipetted onto the coated membrane.

#### Q_m_-sert spheroid harvest

Q_m_-serts were directly transferred from the 96-well plate into the groove of the M-sert, establishing an MQ_m_-sert. The spheroid was carefully pipetted onto the membrane in 150 µL of serum-free DMEM, after which the Q_m_-sert was removed. For both workflows and suspension cells, plates with M-serts were incubated for 30 h to allow cells to invade the Matrigel and migrate to the other side of the membrane. Depending on the analysis, membranes were processed accordingly as described in the following sections.

### Visualization of cells

#### Washing and fixation

After the specified incubation time, cell culture medium was aspirated from M-serts, and they were transferred to another well containing 500 µL DPBS (with Mg^2+^, Ca^2+^), 200 µL was additionally added to the top of the membrane, and the M-serts were washed for 3 min. M-serts were transferred to a well containing 3.7 % formaldehyde in DPBS to fixate cells for 10-15 min. The top part of the membrane was then scraped with a wet cotton swab, after which the M-serts were washed again in DPBS (with Mg^2+^, Ca^2+^) for 2-3 min. If necessary, cells were imaged via fluorescence microscopy. Membranes could be removed, also to facilitate mounting and possible other staining protocols.

#### Crystal violet staining

To visualize cells on the membrane for optical microscopy, cells were stained with 0.3 % crystal violet (CV), which binds to proteins, DNA, and polysaccharides. The CV staining solution was prepared by dissolving CV (0.3 w/v %) in 20 % v/v methanol and vortexing vigorously. After washing and cell fixation, M-serts were transferred to a well containing 500 µL of 0.3 % CV solution and placed on an orbital shaker at 180 rpm for 15 to 20 min, following which the M-sert was dipped in a container containing tap water and then transferred further to another container with ∼500 mL of tap water for 3 min. The samples were dried on lint-free paper towel and could be stored for imaging.

### Microscopy

#### Optical and fluorescence microscopy

Optical microscopy was done with a Zeiss Observer A1 (Carl Zeiss, Germany). Combined optical and fluorescence microscopy was either carried out with a Zeiss Axio Observer Z1 (Carl Zeiss, Germany) or an ECHO Revolve 4K (BICO Group, Sweden). If necessary, images were analyzed and edited using ImageJ 1.53c (NIH, USA). For editing, uneven illumination was reduced using the built-in image calculator process, or brightness and contrast were adjusted.

#### Scanning Electron Microscopy (SEM)

To prepare the membrane and M-sert samples for SEM, samples were mounted on a conductive carbon tape and sputter-coated with gold for 60 s. Samples were imaged using a scanning electron microscope (FEI Quanta 250 FEG, FEI, USA) controlled via xT Microscope Control Software. The images were acquired at an accelerating voltage of 20 kV under soft vacuum (100 Pa) at different magnifications with a large field secondary electron detector. Membrane images pre- and post-sterilization were acquired in high vacuum (< 0.01 Pa).

### Atomic Force Microscopy

Surface scans were performed in tapping mode with a phosphorus-doped silica cantilever in air (k = 3 N/m, f_0_ = 74–90 kHz) at a scan rate of 0.2 Hz with 128 lines per image using a Dimension V atomic force microscope (Bruker Ltd., USA) controlled using Nanoscope software (V.7.3; Bruker Ltd., Billerica, MA, USA). At least three different sample regions, with a scan size of 20 µm², were measured for each condition. NanoScope Analysis (V.1.40; Bruker Ltd.) was used for data analysis and visualization.

### Statistical analysis

All quantitative results in this study were expressed as mean ± standard deviation. Further, statistical analysis was performed using one-way analysis of variance (ANOVA) and Student’s t-test, and F-test for variances. Values of p < 0.05 were considered significantly different: * p < 0.05, ** p < 0.01, n.s. not significant.

### Photography

For taking images of objects, either an iPhone SE 2022 (Apple Inc., USA) or an Alpha 6000 (Sony, Japan) equipped with an interchangeable SELP1650 lens was used.

### Use of artificial intelligence

The use of language editing tools and generative artificial intelligence tools was done according to the Policy on the Use of Generative AI in Research (University of Freiburg, 2024). ChatGPT versions 5.2 and 5.4 were used for image generation (Figure 3B) and language editing; Mistral AI Large 2 (mistralAI, 2024) was used for language editing, both via the OpenWebUI workspace hosted by the University of Freiburg (Germany).

## Results & Discussion

Due to their ability to recapitulate the tumor microenvironment and mimic tumor traits to varying degrees, multicellular or STEM spheroids align with the mandate of EC and FDA for 3DCC to replace animal models. Here, we developed 3D-printed labware for spheroid preparation, carrying out subsequent migration/invasion studies in innovative MQ_m_-serts. Those systems have to yield correct and reproducible results for broad application and can bring down costs in drug discovery.

### Outcomes from M-sert versus commercial control

To ensure a functional system with low margin for errors, emphasis was placed on ergonomic design and the integrity of the membrane bonding in the M-sert. PLA filament was chosen as it is biobased, biodegradable, with an established cytocompatibility profile, and excellent printability in FDM, and furthermore yields prints that are authentic to the CAD model.^23^ The PLA scaffold was designed to be compatible with standard commercial 24-well plates and comparable to commercial inserts, with a holder and two pins for stable attachment, as well as a tightly attached membrane using an epoxy resin (**Figure S1A**). The tightness of the membrane seal was assessed by SEM, where a contiguous layer of epoxy resin was clearly visible between the PLA scaffold and the membrane (**Figure S1B, C**). A further centrifugation test with a common M-sert and a non-porous PET-film control was conducted and showed no sign of leakage (**Figure S1D, E**). Since sterilization is an essential step in cell culture, the membrane was inspected under a microscope before and after sterilization, and no visible alterations were observed (**Figure S2A, B**). Since the assembly of the membrane could potentially transfer pathogens, a mycoplasma test was carried out, which ruled out contamination (**Table S1**). Since epoxy resin was used to attach the membrane, the potential cytotoxicity was evaluated using a standard MTT assay. In comparison with untreated controls, no significant differences in cell viability were observed (**Figure S2C, D**).

To assess the comparative performance of the M-sert to a commercial control, the migration of MDA-MB-231 cells constitutively expressing blue fluorescent protein (BFP) was followed using optical and fluorescence microscopy. The fluorescently labeled cells could be observed without any interference in both systems (**Figure 2A**). Migratory behavior on MDA’s was compared using a standard migration assay protocol using crystal violet (CV) to stain and visualize migrated cells (**Figure 2B**). The number of migrated cells in the control (6351 ± 224) and M-sert (6777 ± 500) was statistically similar (**Figure 2B**). While not statistically significant, a slightly higher number of migrated cells was observed in the M-sert, which could be attributed to ∼3.32% greater surface area of the M-sert membrane.

**Figure 2:**
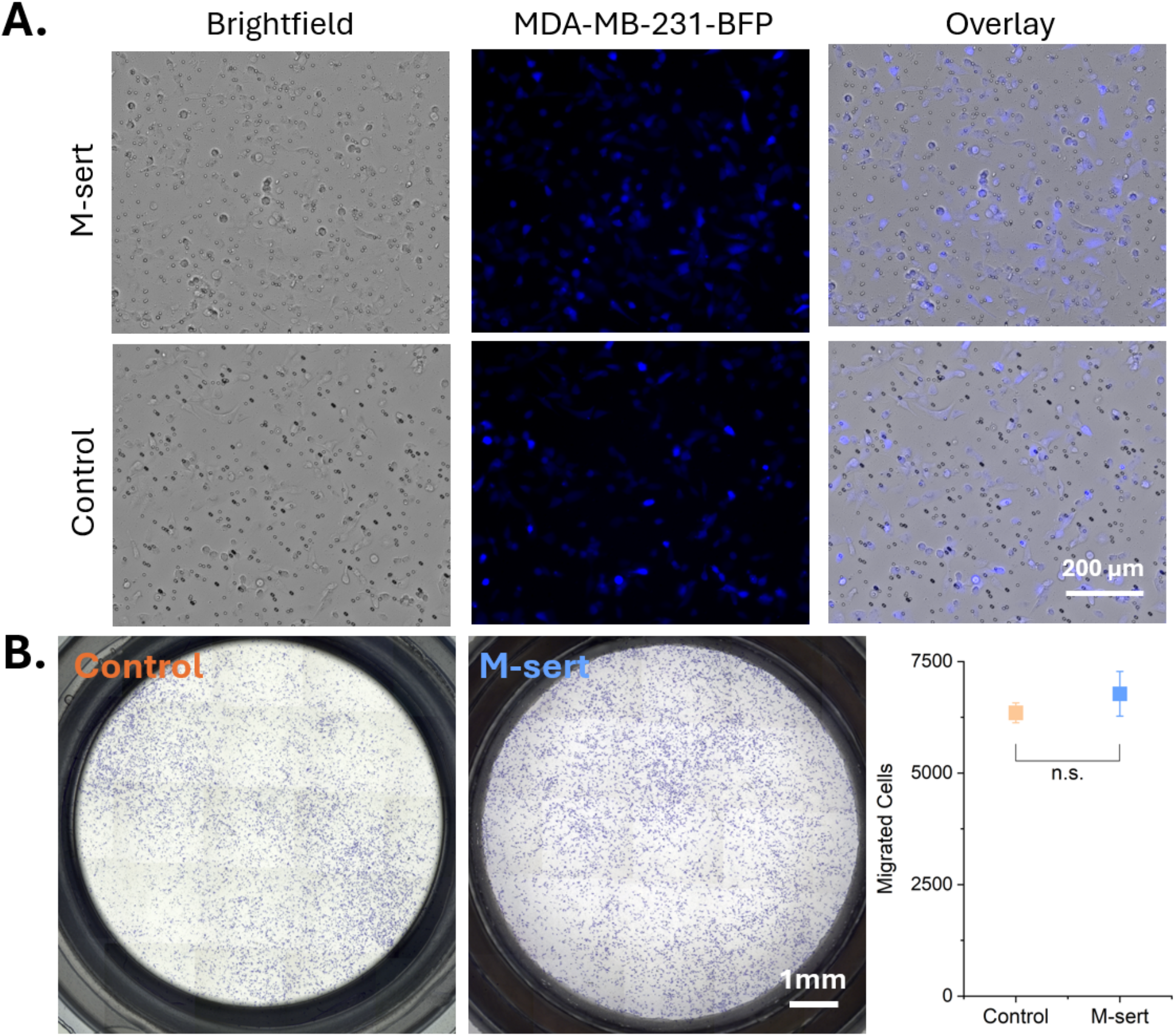
Comparison of M-sert against commercial control reveals no differences. A. Microscopic comparison of MDA-MB-231-BFP cells; scale bar 200 µm. B. CV stain of migrated MDA-MB-231 cells and migration comparison; n = 3 (control), n = 4 (M-sert), two-sample t-test, n.s. not significant.

The absence of cytotoxicity due to the epoxy adhesive in the M-serts is consistent with literature reports that epoxy resins used in dentistry or microfluidics have little to no impact on cell growth.^24–26^ The combination sterilization protocol involving 70% ethanol followed by brief UV exposure had no deleterious effect on the integrity of the PET membrane. It is known that significantly longer UV irradiation of a few hours is needed to affect the integrity of the PET membrane.^27^ The ethanol cleaning step, in addition to disinfecting the surface, might also have aided in the removal of hydrophobic contaminants and positively impacted cell adhesion. Additionally, UV irradiation could increase surface hydrophilicity and improve cell adhesion.^28^ While no discernible differences were observed in membrane wetting behavior, whether the UV treatment step influences cell migration remains to be investigated.

The M-sert design deliberately relied on the same principles as commercial products to ensure direct comparison and adoption by the research community. To further standardize protocols and improve throughput efficiency, researchers have engineered membranes that prevent fluorescent bleed-over from the upper membrane surface. This design eliminates the need to detach the cells from the top compartment, permitting direct imaging and analysis.^29^ Together with automated cell counting and other analyses developed for migration and invasion assays, this could further improve the M-sert workflow.^30^

### Design Iteration of the MQ_m_-sert

Migration is a prerequisite for invasion, and invasiveness is a key determinant of metastatic potential of cancer cells. Therefore, the invasion of cells from multicellular STEM spheroids composed of mesenchymal cells, triple negative breast cancer cells (MDA-MB-231) and microvascular endothelial cells was studied. As noted in the introduction and Figure 1, a traditional two-step handling of the spheroid for a membrane migration/invasion assay might dissociate/disrupt the spheroid, thus introducing variability and skewing the results (**Figure 3**). When handling and harvesting spheroids prepared using a conventional commercial handling drop system, detached cells and cellular aggregates were observed (**Figure 3**), confirming our reasoning and emphasizing the need to improve the two-step handling process.

**Figure 3:**
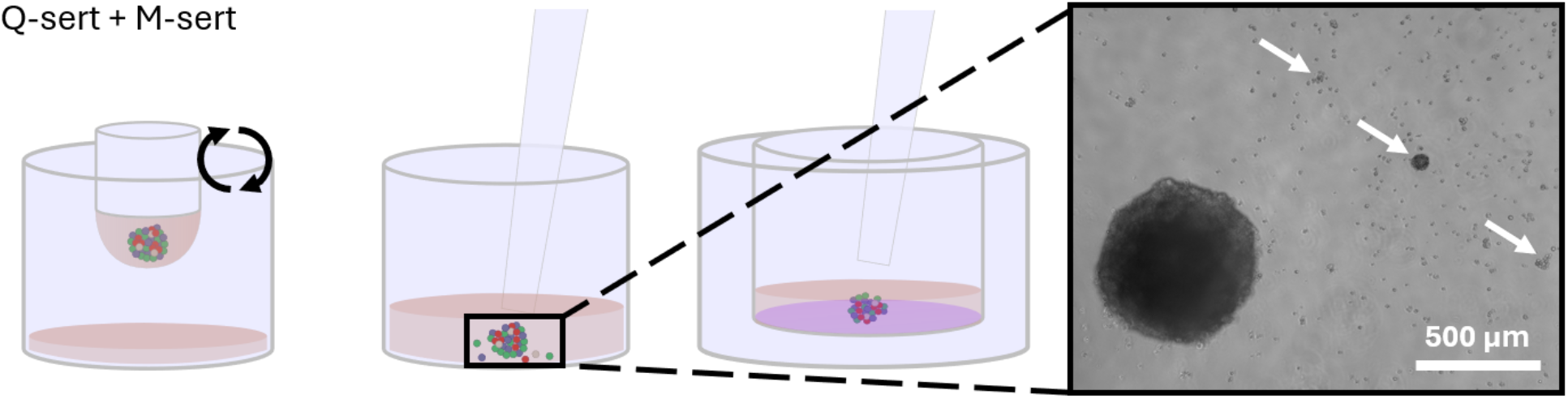
Conventional spheroid harvesting protocol disrupts spheroids. Handling and transfer of a STEM spheroid in a traditional workflow involving a hanging drop culture followed by transfer to a migration membrane (Q-sert and M-sert) resulted in detached cells and cell aggregates, indicated by white arrows, which are easily discernible in conventional microscopy (scale bar 500 µm).

To overcome this issue of spheroid disintegration, the workflow was reimagined to skip the spheroid handling step. Towards this objective, the Q-sert and M-sert were redesigned to facilitate a single-step transfer, resulting in an MQ_m_-sert device (**Figure 4**). The redesign involved multiple iterations to meet the identified requirements. In the 1^st^ generation MQ-serts, a single-part construction led to excessive evaporation of the hanging drop in 24-well plates (**Figure 4A**). Additionally, the lid contact at the top of the MQ-sert caused loss of culture medium due to capillary forces. To address these issues, in the 2^nd^ generation design, the device was split into two components: the M-sert featuring an edge groove and the Q_m_-sert as a detachable insert (**Figure 4A**). However, lid interference persisted, prompting a third iteration in which the groove was deepened, and the Q_m_-sert was modified with a counterpiece to the groove’s fillet (i.e., the rounded corner transitioning from the edge to the surface) for a secure fit (**Figure 3A**, **Figure S3A**).

**Figure 4:**
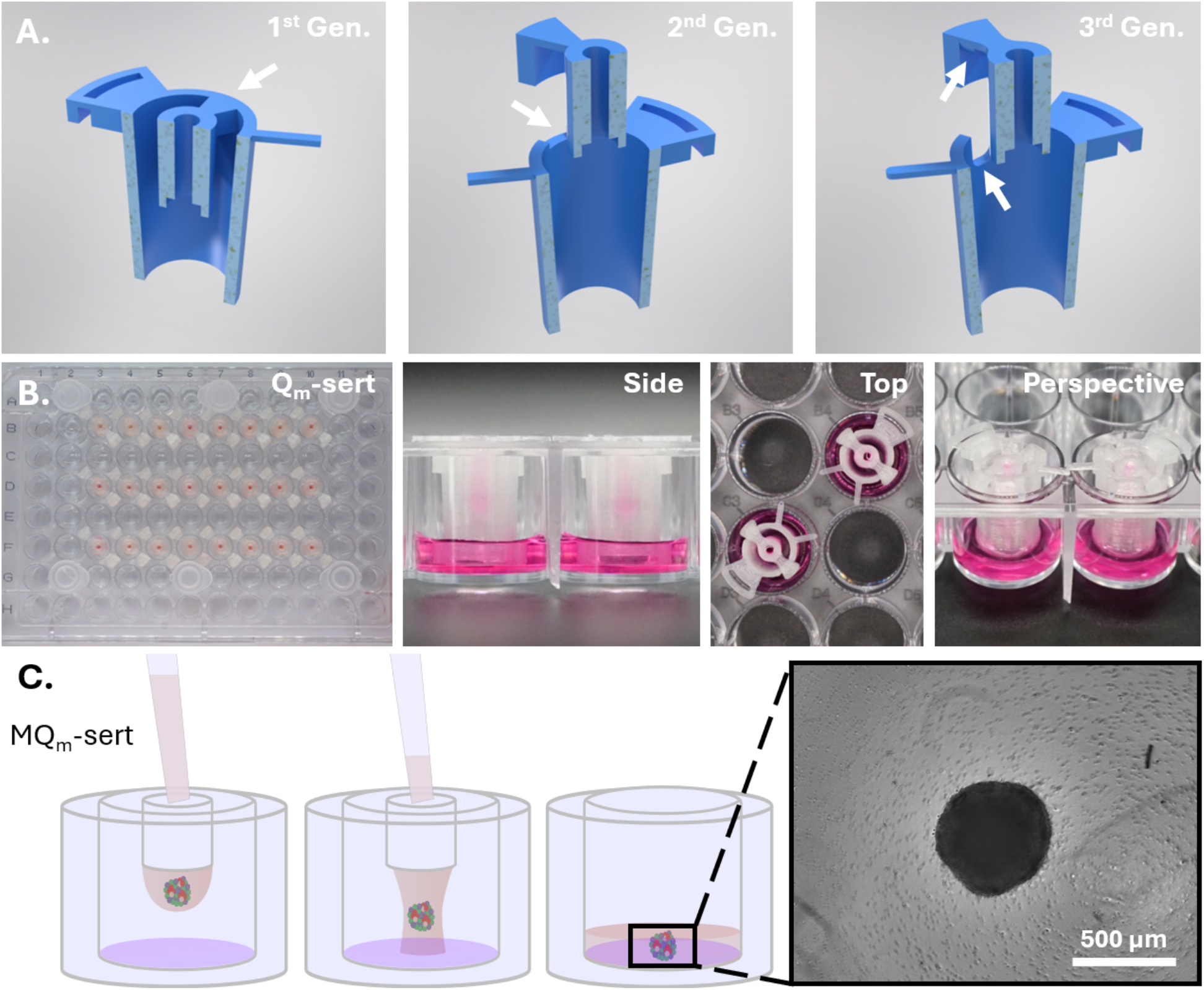
Design evolution of the MQ_m_-sert allows gentle spheroid transfer onto the membrane. A. Design evolution and improvement of the MQ_m_-sert in three iterations. B. Photographs of Q_m_-sert spheroid culture and the assembled MQ_m_-sert. C. STEM spheroid transfer procedure in a combined setup (MQ_m_-sert) with spheroid microscopy after harvest; scale bar 500 µm.

The workflow of the manufacturing and application of MQ_m_-serts is shown in **Figure S3B**: Both components that make up the MQ_m_-serts were manufactured by FFF, processed (cleaning, membrane attachment), and sterilized, making them ready for use in cell culture applications (**Figures S3B, 4B**). Following spheroid culture, the Q_m_-sert was transferred to the M-sert, enabling the careful deposition of the spheroid-containing droplet onto the membrane (**Figures 4C, S3B**). This integrated workflow eliminated the limitations of conventional two-step protocols, as spheroid integrity was preserved, as confirmed by optical microscopy (**Figure 4C**).

For the development of innovative labware, e.g., MQ_m_-serts, the FFF is highly suitable as it supports an iterative process as shown here, where prototypes (i.e., generations 1-3, **Figures 4A)** were rapidly fabricated, allowing design optimization to fit application criteria.^17, 19^ This study demonstrates the value of cross-disciplinary incorporation, particularly from researchers in synthetic biology, engineering, and product development, where the “design-build-test-learn” cycle has proven to be an effective iterative strategy for refining successive generations of functional devices.^31^

### Comparison of operational workflow: Q- and M-sert vs. MQ_m_-sert and invasion behavior of STEMs in MQ_m_-sert

To quantitatively prove that the observed differences are reflected in the actual behavior of the invasive phenotype of STEMs, the Q-sert + M-sert workflow (two-step) was compared to our innovative MQ_m_-sert workflow (one step). To minimize inter-experimental variability, the comparison was performed using cells from the same batch for both workflows and in parallel. Both setups yielded the same STEM spheroid size (Q-sert 524 ± 17 µm; Q_m_-sert 518 ± 26 µm) and cellular organization (**Figure S4**).

To analyze the invasion capacity of the single cell types of a STEM spheroid, the invasion behavior was first characterized for MDA-MB-231, MCF7, HPMECs, and MSCs by themselves as 2DCC (**Figure S5**). Unlike cellular migration, cell invasion requires the degradation of ECM to facilitate movement into a specific location or direction. Hence, the membrane was coated with Matrigel, an ECM extract which is the gold standard for cell invasion experiments in cancer biology.^32–34^ Among the tested cell types, only MDA-MB-231 cells exhibited highly invasive capacity, whereas MSCs presented significantly fewer invasive cells. (**Figure S5**). Invasive cells from STEM spheroids were quantified (**Figure 5**) for workflow comparison. CV-stained cells (MDA-MB-231) as well as fluorescent MSCs were imaged via microscopy and counted. Surprisingly, no differences were observed for MDA-MB-231 cells from both workflows, with 1655 ± 609 (Q) and 1623 ± 378 (Q_m_) cells (**Figure 5**). However, we observed a significantly different number of invasive MSCs, with 47 ± 15 (Q) and 64 ±11 (Q_m_) cells, highlighting the influence of the mentioned workflows on the experimental outcome. Additionally, the difference could also be quantified by calculating the ratio of MDA-MB-231 to MSC, which could give information about the quantitative relationship between breast cancer and stromal cell invasion (Q ∼ 38, Q_m_ ∼ 26; **Figure 5**).

**Figure 5:**
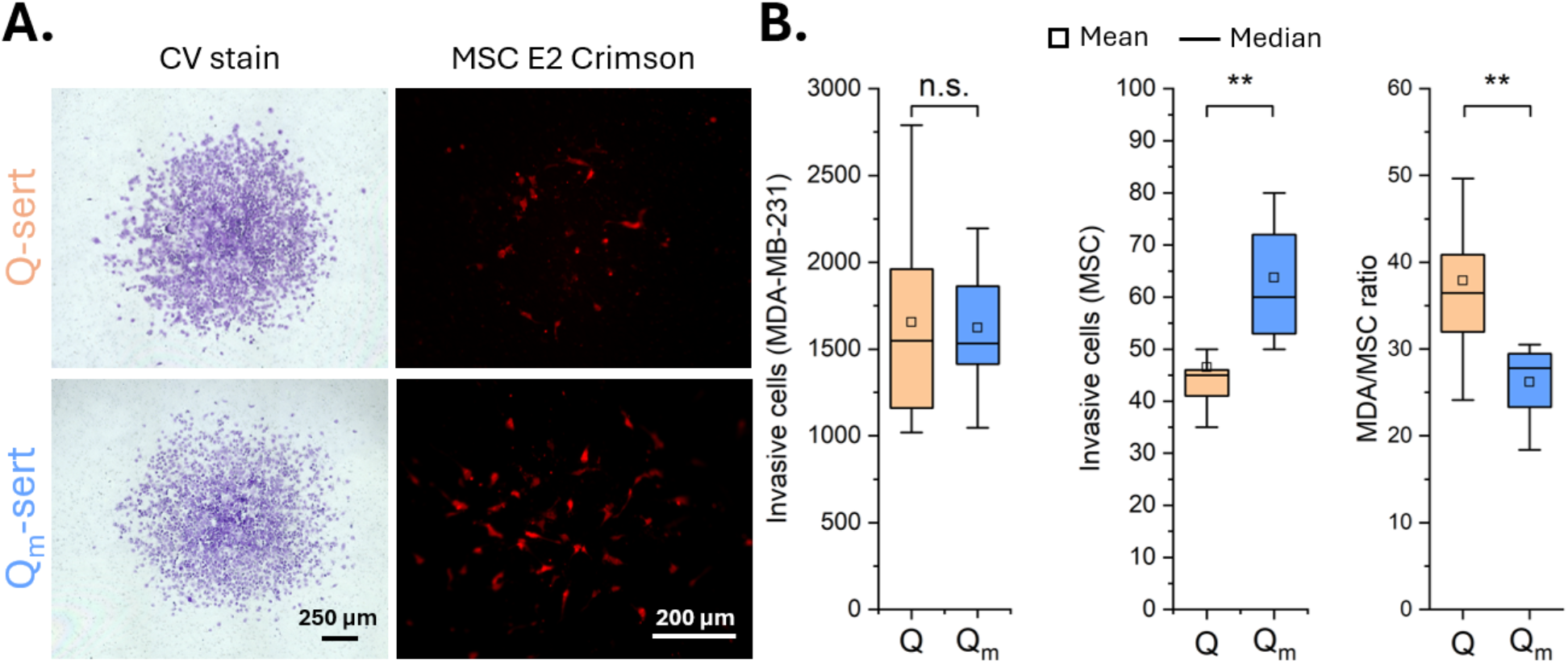
Differences in MSC invasion behavior introduced by the invasion assay. A. Exemplary brightfield microscopy color image of CV-stained cells (left) and fluorescent microscopy image of human MSCs (right); scale bar 250 µm (left), 200 µm (right). B. Quantification of the invasive MDA-MB-231 cells and MSCs and their ratio (Q-sert: n = 12, Q_m_-sert: n = 9). Two-sample t-test, n.s. not significant, **p <0.01.

In contrast to our setup and workflow, the general setup of the membrane invasion assay has not changed for years, nor have any modifications been made. However, other assays, namely cells or spheroids embedded in an ECM, such as collagen, have been developed and used to study invasion. This offers a clear advantage over Matrigel-coated membranes, as embedding mimics the three-dimensional physiological environment provided by the extracellular matrix.^35–38^ This physiological environment can also be recreated on the MQ_m_-sert by covering harvested spheroids with ECM. While those prior studies similarly harvested spheroids, they either failed to address the spheroid integrity challenges described herein or required specialized, non-standard equipment (e.g., custom-fabricated plates) that exceeds the capabilities of conventional laboratory setups.^35–37, 39^ Reports on the importance of spheroid handling are rather rare,^40^ and in this study, one can speculate that disruption of the spheroid leads to disintegration of the ECM and detachment of MSCs, which then do not contribute to the invasive cells. Interestingly, a disruption to the 3D architecture of a spheroid is connected to a more malignant phenotype for colorectal cancer,^41^ and therefore, specific molecular differences could be probed in Q vs. Q_m_ workflows in the future, although this might not lead to differences given the identical culture conditions. The invasive capacity of MDA-MB-231 cells and MSCs, and their cellular interplay, e.g., for the degradation of ECM by metalloproteinases, follows already reported studies.^42–44^ As part of the STEM spheroid, MSCs play multiple roles in the tumor microenvironment, and can be seen as the precursor of cancer-associated fibroblasts promoting tumorigenesis, which is why correct reporting with the MQ_m_-serts is important.^45^

### In-depth data analysis of operational workflow and cost comparison

As already pointed out earlier, the data quality of 3D models is vital to draw meaningful and robust conclusions. Here, we aimed to analyze the data generated in detail to gain quantitative insights. We measured the dimensions of the stained cell mass after invasion on the membranes, considering the invasive cell mass as elliptical or circular. The expansions of the cell mass in x-and y-directions, their area, as well as the invasive cells per mm², were examined to gain insights into the geometry of the invading cell mass from the differently prepared workflows Q vs. Q_m_ (**Figure 6**).

**Figure 6:**
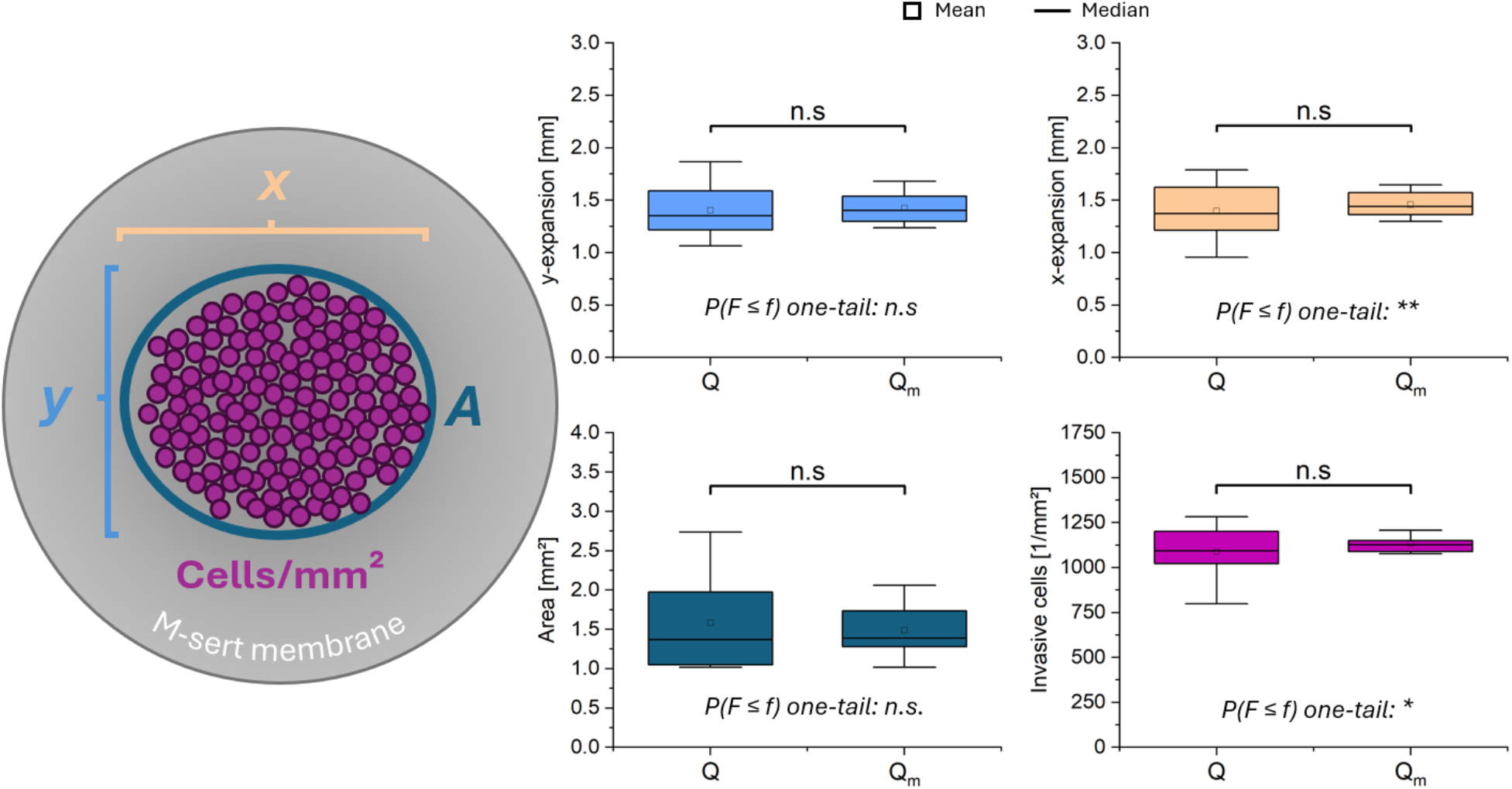
The Q_m_-sert workflow delivers less data variance. Dimensions of invasive cell mass were determined on the membranes for x- and y-dimensions, the covered area, and cells/mm². Two-sample t-test and two-sample F-test for variances, n = 12 (Q-sert), n = 9 (Q_m_-sert), n.s. not significant, *p < 0.05, **p < 0.01.

Upon comparing the four listed criteria, no significant differences between the two workflows were observed in a t-test (**Figure 6**, **Table 2**). However, the data are more broadly distributed in the Q-sert workflow, as confirmed by an F-test for variances indicating significantly different variances for x-expansion and invasive cells per mm² (**Figure 6**, **Table 2**). The elevated area in the Q-workflow, as well as fewer cells per mm², can be attributed to and seen as a confirmation for the disintegration of the spheroids, becoming less compact, and therefore likely to spread more on the membrane.

**Table 2:**
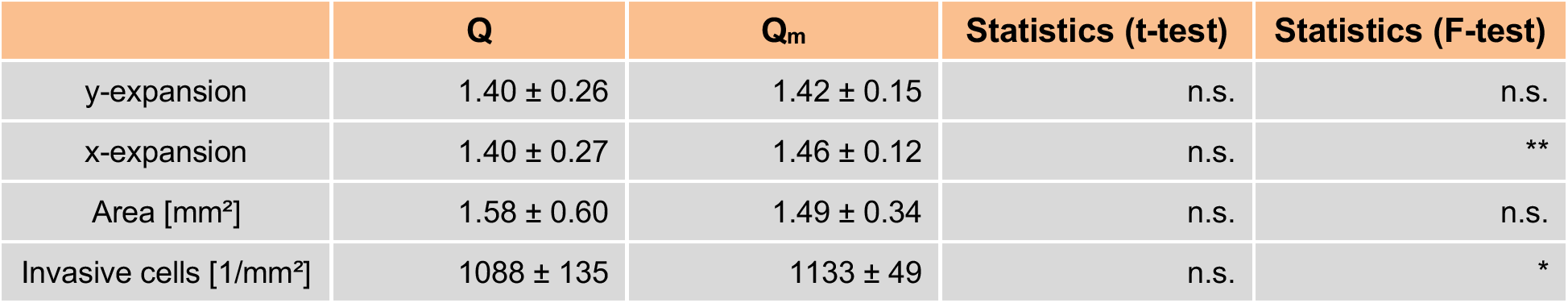
Data variance in the Q_m_-sert workflow. Variance was assessed on data shown in Figure 5. Two-sample t-test and two-sample F-test for variances, n = 12 (Q-sert), n = 9 (Q_m_-sert), n.s. not significant, *p < 0.05, **p < 0.01.

|  | Q | Q<sub>m</sub> | Statistics (t-test) | Statistics (F-test) |
| --- | --- | --- | --- | --- |
| y-expansion | 1.40 ± 0.26 | 1.42 ± 0.15 | n.s. | n.s. |
| x-expansion | 1.40 ± 0.27 | 1.46 ± 0.12 | n.s. | ** |
| Area [mm<sup>2</sup>] | 1.58 ± 0.60 | 1.49 ± 0.34 | n.s. | n.s. |
| Invasive cells [1/mm<sup>2</sup>] | 1088 ± 135 | 1133 ± 49 | n.s. | * |

Other researchers have also highlighted the need for reproducibility and standardization in spheroid preparation^11, 46–48^, and our results address these demands for reproducible and standardized spheroid handling in cancer research, for example, for drug screening. Finally, because fabrication of bespoke labware has to be price-wise competitive with commercially available products, we conducted a comparison just for the M-sert and a commercial standard, considering the materials’ costs as well as the labor going into fabrication. The price per piece was € 9.12 (for the commercial standard) vs. € 2.24 (M-sert), which amounts to a ∼ 75 % saving. The competitive price, especially for highly specific labware and small quantities, but also the democratization of science for lower-income countries and laboratories with fewer resources, are also reported in other studies.^16–18^

## Conclusions

The MQ_m_-sert platform represents a significant advancement in 3D cell culture technology, addressing key limitations of traditional spheroid-based migration and invasion assays. By integrating hanging-drop spheroid culture with membrane-based analysis in a single workflow, this 3D-printed, open-source system eliminates disruptive spheroid transfer steps, preserving structural integrity and reducing experimental variability. Its cost-effective fabrication using biobased PLA and compatibility with standard 24-well plates enhance accessibility and sustainability, aligning with regulatory mandates to replace animal testing with 3D cell culture models. Beyond technical improvements, the MQ_m_-sert enables standardized, scalable investigations of tumor-stroma interactions, offering a relevant tool for cancer research and drug screening.

## Acknowledgements

The authors kindly thank Prof. Andrea Barbero for providing the marrow-derived mesenchymal stroma cells. Thanks to Dr. Nicole Gensch from the CIBSS/BIOSS Toolbox, (University of Freiburg, Germany) for providing cells and plasmids. Tobias Butelmann acknowledges Wissenschaftliche Gesellschaft in Freiburg im Breisgau and Dr. Leo-Ricker-Stiftung. The authors would like to thank Vincent Ahmadi for acquiring SEM images.

## Author Contributions

Conceptualization, TB and VPS; methodology, TB; formal analysis, TB and TN; investigation, TB and TN; resources, VPS; data curation, TB and TN; writing—original draft preparation, TB; writing—review and editing, all authors; visualization, TB; supervision, TB and VPS; project administration, TB and VPS; funding acquisition, TB and VPS. All authors have read and agreed to the published version of the manuscript.

## Supplementary Materials

Supplementary Materials, Figures S1-S5 and Table S1, are attached to this file.

## Supplementary information

**Figure S1:**
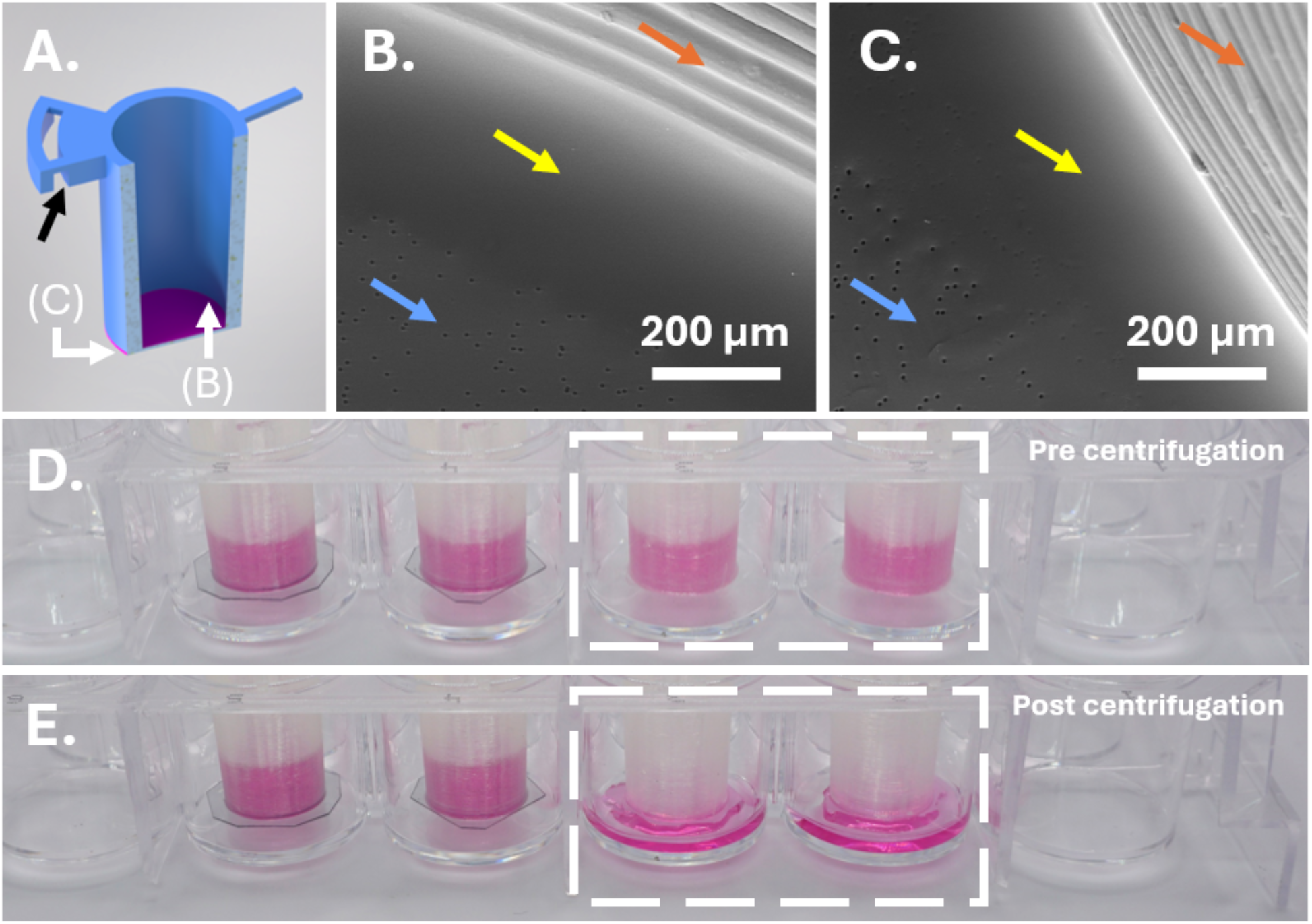
Leakproofness of the M-sert. A. Rendered M-sert design as cross section with attached membrane (purple); black arrow indicating holder, white arrows indicating SEM micrograph sites. B. SEM micrograph of PLA scaffold and PET membrane (inside); scale bar 200 µm. C. SEM micrograph of PLA scaffold and PET membrane (outside); scale bar 200 µm. D. Pre-centrifugation: M-serts filled with 200 µL DMEM prior to centrifugation; dashed rectangle indicating membrane samples, left of it with tight PET film. E. Post centrifugation: M-serts filled with 200 µL DMEM prior to centrifugation; dashed rectangle indicating membrane samples, left of it with tight PET film.

**Figure S2:**
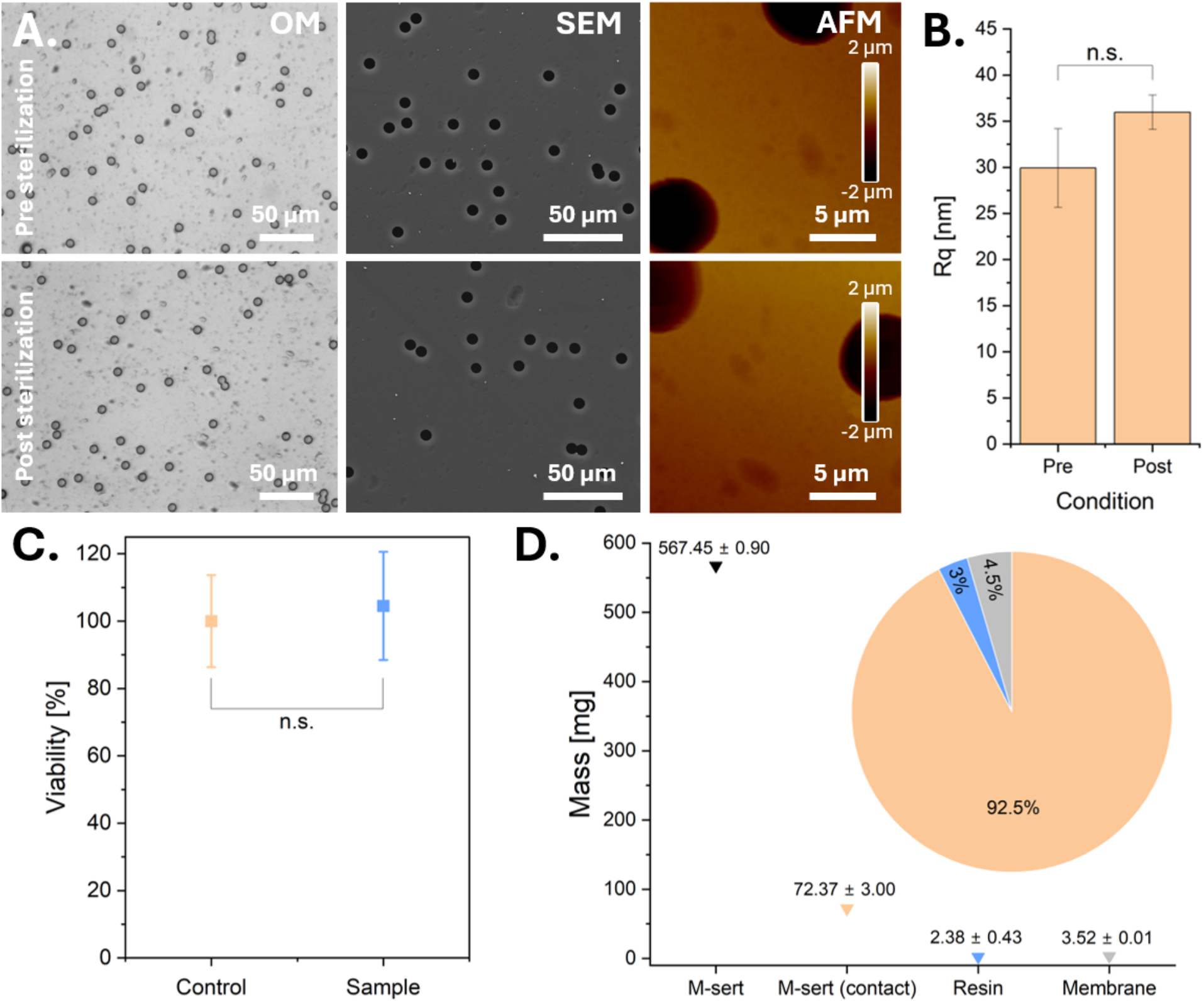
The fabrication and sterilization do not influence membrane integrity or cell viability. A. Microscopic investigation of the membrane before and after sterilization; scale bar (OM) 50 µm, (SEM) 50 µm, (AFM) 5 µm, z-scale 2 µm. B. Surface roughness (Rq) of the membrane before and after sterilization, n = 3, two sample t-test, n.s. not significant. C. MTT assay to assess cell viability after immersing the M-sert in complete cell culture medium compared to an untreated control; n = 5, two sample t-test, n.s. not significant. D. Mass touched by complete cell culture medium during M-sert immersion, fractions indicated in pie chart; n = 3.

**Table S1:** Absence of mycoplasma after sterilization. Test carried out by Eurofins Genomics, n = 3.

| Identification |  | Results |  |  |
| --- | --- | --- | --- | --- |
| Sample No. | Barcode | PCR inhibition | Mycoplasma | Summary |
| 1 | MP00024715 | Absent | Absent | Clean |
| 2 | MP00024717 | Absent | Absent | Clean |
| 3 | MP00024718 | Absent | Absent | Clean |

**Figure S3:**
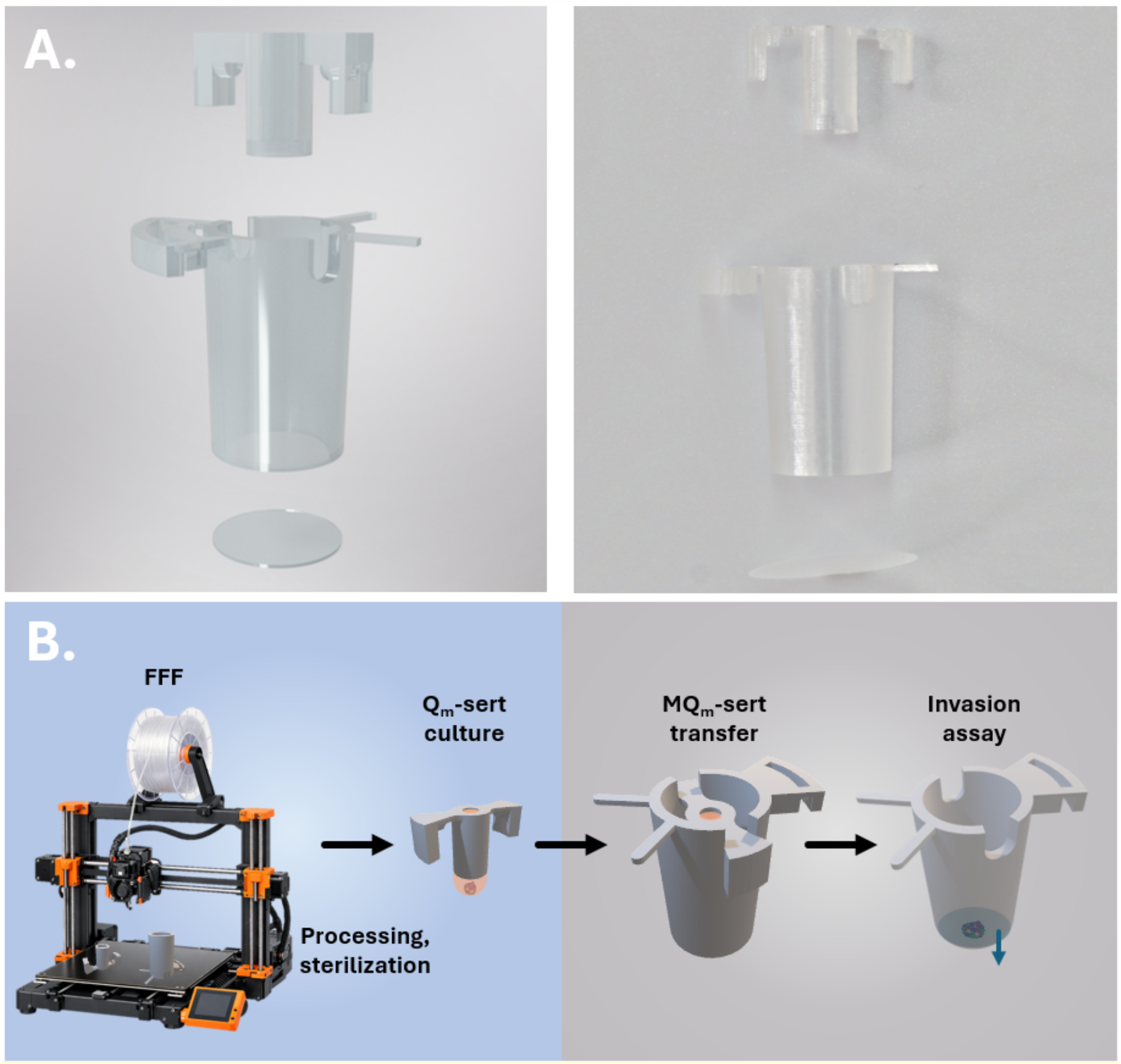
Final MQ_m_-sert design and workflow. A. Rendered image and photographs of Q_m_-sert, M-sert and PET membrane of the third generation. B. Fabrication, processing, spheroid culture, and invasion assay workflow with MQ_m_-serts.

**Figure S4:**
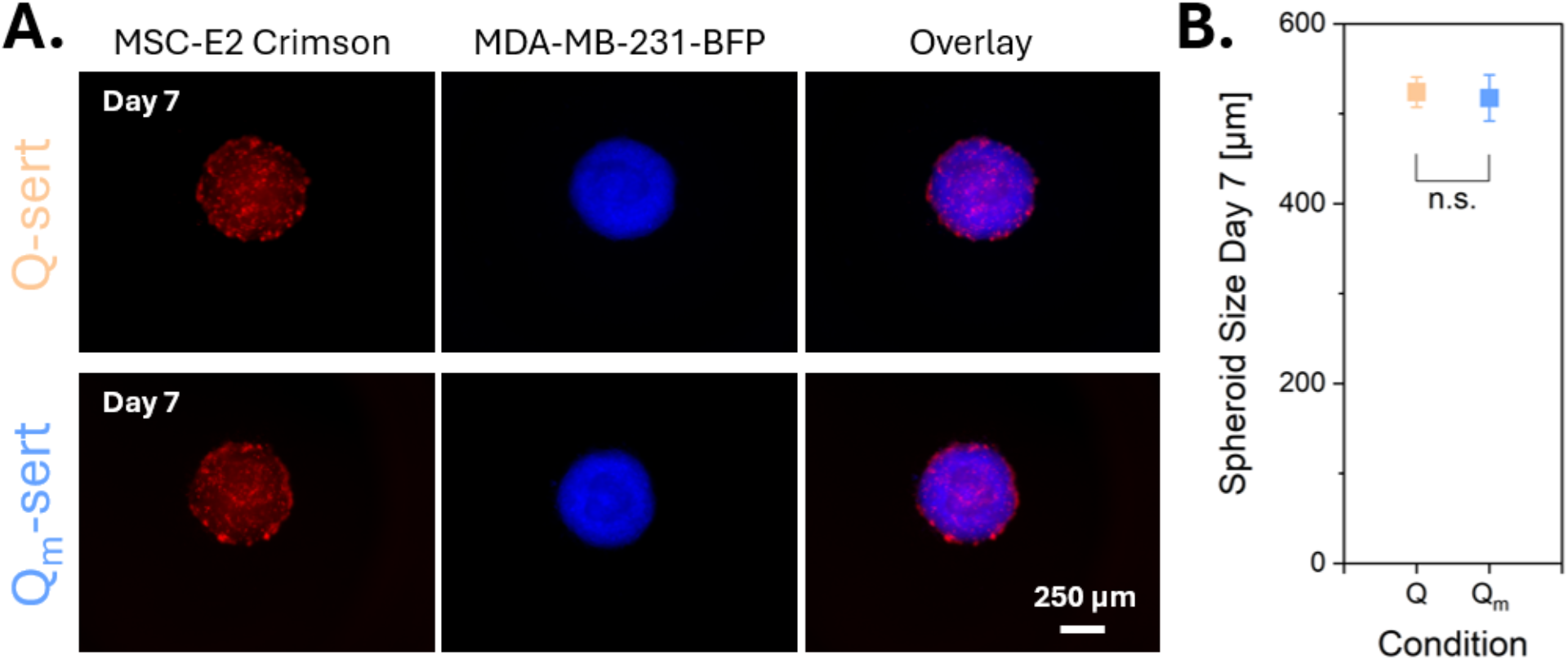
STEM spheroids cultured in Q- and Q_m_-serts show the same size and cellular organization. A. Fluorescence microscopy of STEM spheroids in Q- and Q_m_-serts on day 7; scale bar 250 µm. B. Spheroid size analysis of spheroids cultured in Q- and Q_m_-serts on day 7; n = 10 (Q_m_), n = 20 (Q), two sample t-test, n.s. not significant.

**Figure S5:**
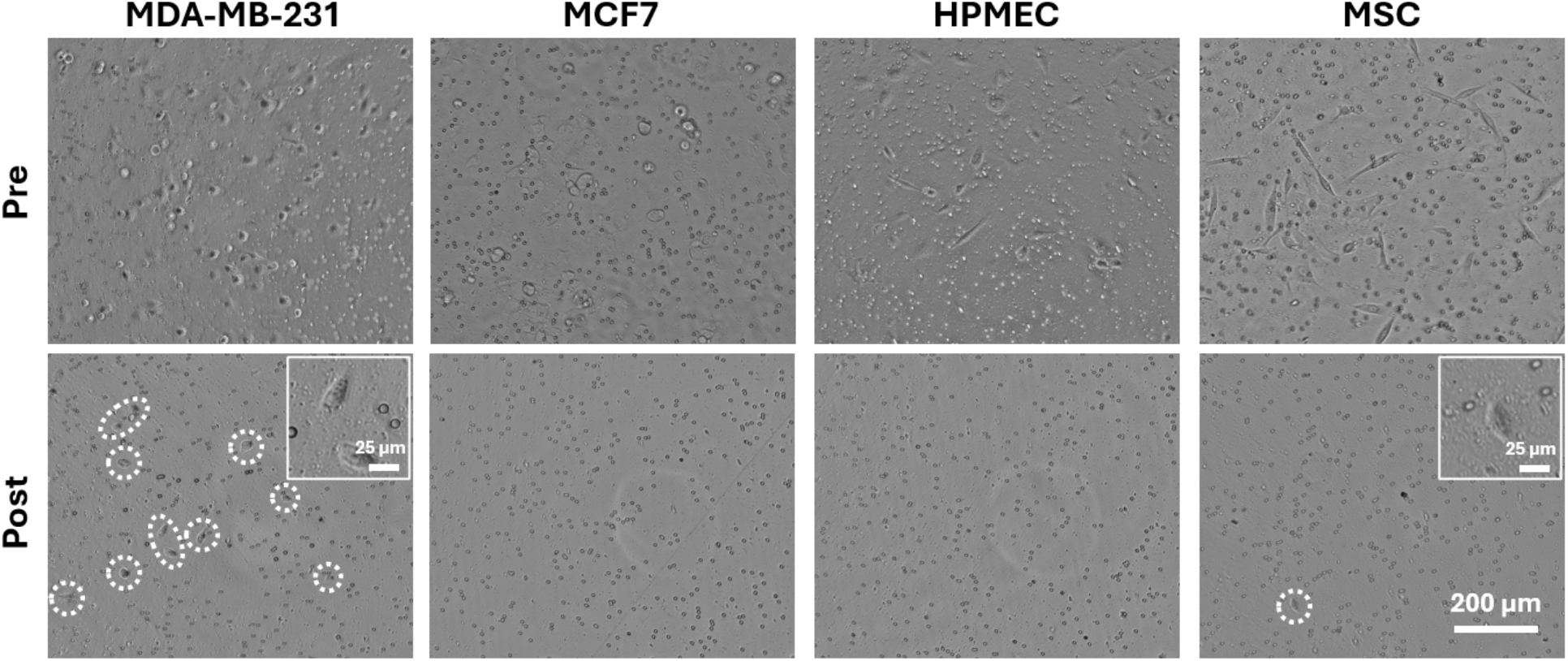
Invasion was present in MDA-MB-231 cells and less pronounced in MSCs. Optical microscopy of single cell types in an invasion assay before and after removal of non-invasive cells from top of the membrane; dashed circles indicating invaded cells, scale bar 200 µm. Integrated magnified window in post condition showing cells, scale bar 25 µm.

